# The false mussels (*Mytilopsis leucophaeata*) can be mechanical carriers of the shrimp microsporidian *Enterocytozoon hepatopenaei* (EHP)

**DOI:** 10.1101/2021.04.23.441221

**Authors:** Natthinee Munkongwongsiri, Orawan Thepmanee, Kanokwan Lertsiri, Rapeepun Vanichviriyakit, Ornchuma Itsathitphaisarn, Kallaya Sritunyalucksana

**Affiliations:** Aquatic Animal Health Research Team (AQHT), National Center for Genetic Engineering and Biotechnology (BIOTEC), National Science and Technology Development Agency (NSTDA), Yothi office, Rama VI Rd., Bangkok 10400, Thailand; Center of Excellence for Shrimp Molecular Biology and Biotechnology (Centex Shrimp), Faculty of Science, Mahidol University, Rama VI Rd., Bangkok 10400, Thailand; Department of Biochemistry, Faculty of Science, Mahidol University, Bangkok 10400, Thailand; Department of Anatomy, Faculty of Science, Mahidol University, Bangkok 10400, Thailand

## Abstract

*Enterocytozoon hepatopenaei* (EHP) is an obligate intracellular parasite causing hepatopancreatic microsporidiosis (HPM) in cultivated shrimp in Asian countries. One strategy to control EHP is to identify and eliminate biological reservoir(s) in shrimp ponds. Several marine and brackish-water organisms, including false mussels (*Mytilopsis*) have been reported to test positive for EHP using polymerase chain reaction (PCR) technology. Thus, we tested samples of commonly found Thai false mussel *Mytilopsis leucophaeata* from 6 EHP-infected shrimp ponds by PCR for the presence of EHP using the spore wall protein (SWP) gene primers. The mussel samples from all 6 ponds were positive. Subsequent bioassays carried out using naïve mussels cohabitated with EHP-infected shrimp gave 100% SWP-PCR positive mussels at 20 days. One batch of such PCR-positive mussels was transferred for cohabitation with naïve shrimp and gave 37.5% EHP-positive shrimp within 10 days. Tissue analysis of the EHP-PCR-positive mussels using light microscopy, *in situ* hybridization analysis for the SWP gene and electron microscopy did not confirm EHP infection. In summary, we obtained no evidence that *Mytilopsis leucophaeata* was itself infected with EHP. However, it was apparently capable carrying infectious spores for some period after ingestion and serving as a mechanical or passive carrier. The results support previous reports warning of the danger of feeding living or fresh bivalves to broodstock shrimp in hatcheries or shrimp in rearing ponds without prior heating or freezing.

## INTRODUCTION

*Enterocytozoon hepatopenaei* (EHP) is an obligate, intracellular parasite that currently threatens shrimp culture in several Asian countries (Ha *et al*., 2010; Tangprasittipap *et al*., 2013; Rajendran *et al*., 2016, Shen *et al*., 2019). EHP was first discovered in Thailand in *Penaeus monodon* in 2004 (Chayaburakul, *et al*., 2004) and later described in detail and named in 2005 and 2009 (Tourtip, 2005; Tourtip, *et al*., 2009). Infections with similar morphology were previously reported in *P. monodon* from Malaysia {Anderson, 1989 #8332} and in *P. japonicus* in Australia {Hudson, 2001 #8310}. Perhaps the widespread adoption of exotic *P. vannamei* as the species of choice for cultivation in Asia around 2004 and the subsequent rapid expansion in production together with lack of attention to it made the shrimp cultivation industry more vulnerable to this pathogen. A recent review (Chaijarasphong et al., 2020) of EHP summarizes the current status of research on EHP.

Fortunately, the specific pathogen free (SPF) breeding stocks of *P. vannamei* were mostly developed prior to its introduction to Asia, and it is possible that it does not occur naturally in its native environment of *P. vannamei* in the western Pacific Ocean. Thus, it was not carried by these stocks and it is now possible for all prudent shrimp farmers to avoid stocking of EHP-infected post-larvae (PL). However, there is evidence that widespread infections can occur via contamination of SPF broodstock in shrimp hatcheries or their offspring PL when stocked in shrimp ponds. In these cases, the infections must originate from natural carriers of EHP. Unfortunately, little is still known regarding the potential carriers, even though a number of marine animals used for broodstock feed have tested positive for EHP (Chaijarasphong et al., 2020). Without detailed tissue analysis and bioassay it is not possible to distinguish passive (mechanical) carriers from infected carriers, with the latter being the most likely to constitute a stable environmental reservoir.

In this study, we focused on examining the possibility that *Mytilopsis leucophaeata* might be an infected carrier of EHP. A preliminary examination was carried out for the prevalence of growout ponds that contained shrimp positive for severe EHP infections (by PCR analysis). Those ponds were found the mussel, *M. leucophaeata* positive for EHP by PCR too. Subsequent laboratory tests were carried out using EHP-PCR negative *M. leucophaeata* cohabitated with EHP-infected shrimp to test for EHP transmission from shrimp to mussels. Transmission was successful and the resulting infected mussels were divided into two groups. One group was used directly for cohabitation with naïve shrimp to determine ability of the mussels to transmit EHP to shrimp. Another group was used to carry out tissue analysis by histology, *in situ* hybridization analysis and transmission electron microscopy (TEM) to determine whether or not the EHP-positive mussels were actually infected with EHP.

## MATERIALS AND METHODS

### Shrimp welfare

This work followed Thailand’s laws for ethical animal care under the Animal for Scientific Purposes ACT, B.E. 2558 (A.D. 2015) under project approval number BT-Animal 12/2563.

### Selection of sampled ponds

Altogether 6 shrimp grow-out ponds from 3 districts (Na Yai Arm, Tha Mai and Laem Singha) in Chantaburi province, Thailand were selected. The farmer owners complained that their shrimp were continually infected with EHP. From each pond, samples of both shrimp (*P. vannamei*) and false mussels (*M. leucophaeata*) were collected.

### Preparation of shrimp and mussel tissue samples for PCR analysis

For shrimp, hepatopancreas (HP) from 10 shrimp from each pond was dissected and individually homogenized in 1 ml DNA lysis buffer (buffer (50 mM Tris pH 9, 0.1M EDTA pH 8, 50 mM NaCl, 2% SDS) containing 100 μg/ml of proteinase K in preparation for DNA extraction. Mussels were thoroughly washed externally with clean tap water 3 times before internal tissue were homogenized in the same DNA lysis buffer used for shrimp.

### DNA extraction and PCR detection methods

Samples in DNA lysis buffer were subjected to DNA extraction using a QIAamp® DNA Mini Kits (Qiagen). DNA extracts were subjected to quantification by spectrophotometer (Nano drop 200c, Thermo scientific) prior to use for PCR procedures.

For EHP detection, DNA extracts were used as templates for the EHP-SWP nested PCR method (Jareonlak *et al*., 2016). The primers are shown in Table 1. The PCR mixtures for first step and nested PCR step (12.5 μl) contained 1X OneTaq Hot Start Master Mix (NEB) and 0.2 μM of each primer. The template for the first PCR contained 100 ng of DNA template while that for the nested PCR step consisted of 1 μl of the final reaction solution from the first PCR step. Thereafter, amplicons were analyzed by 1.5% agarose gel electrophoresis with ethidium bromide staining and using a DNA ladder marker (2 log, 100 bp, or 1 kb DNA ladder from New England Biolabs, USA). Expected PCR product sizes of the first step and nested PCR step were 514 and 148 bp, respectively.

**Table 1:**
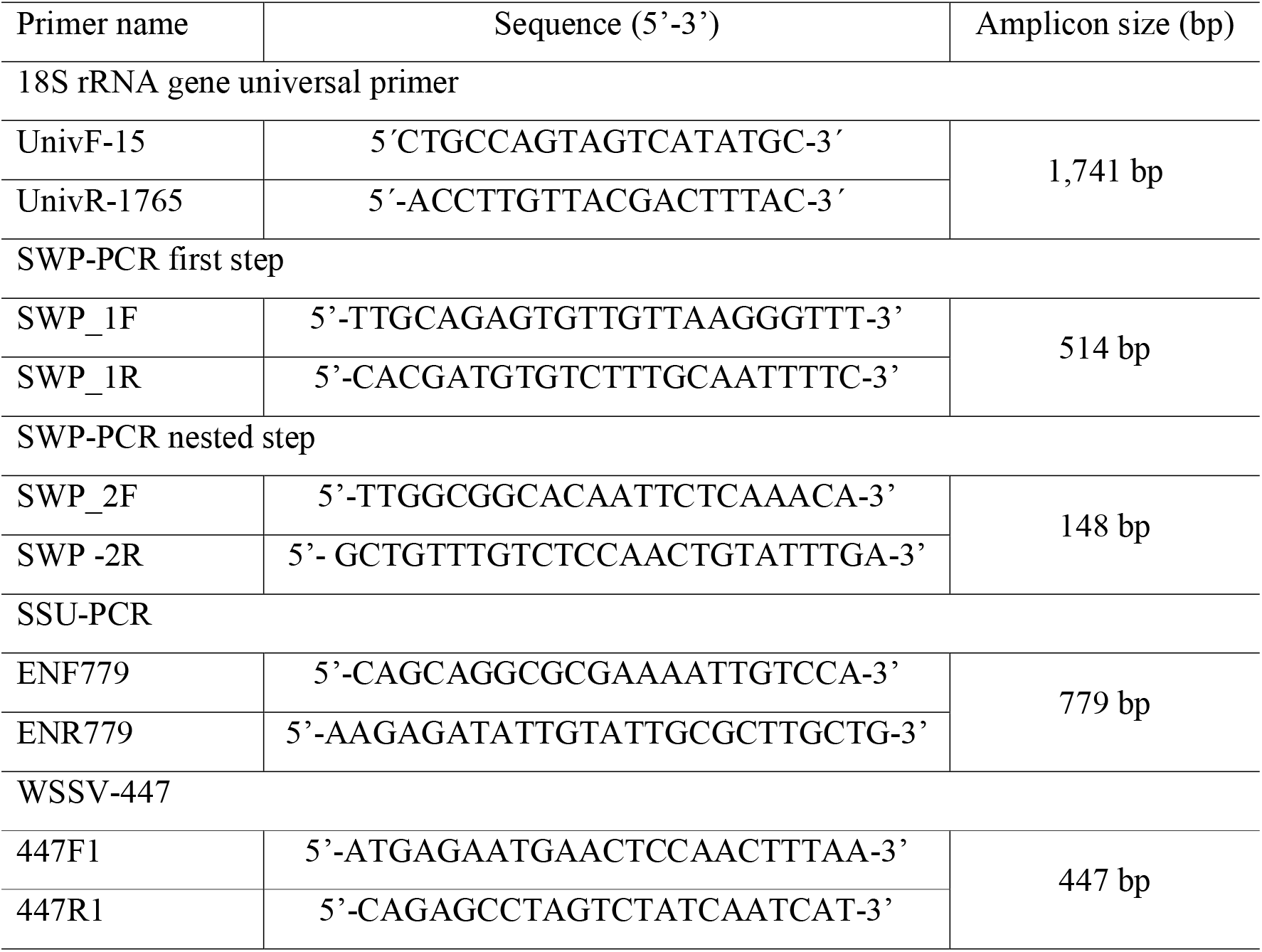
PCR primers used in this study

### Stocks of naïve shrimp and EHP infected shrimp

Naïve *P. vannamei* (size 2-3 g fresh weight, n=40) were transported from a shrimp farm in Chacheongsao province, Thailand and checked to be EHP-free by PCR. While EHP infected *P. vannamei* were taken from a commercial shrimp farm (size 8-10 g fresh weight, n=50) in Chantaburi province, Thailand. Shrimp were acclimatized for 5 days in 500 liter-plastic tanks containing 350 liters of 20 ppt artificial seawater with dissolved oxygen at 5 ppm, pH 7.8, alkalinity 130-150 mg/l and daily feeding at 3% of body weight.

### Collection, screening and identification of Mytilopsis sp

Three hundred mussels (*M. leucophaeata*, Fig. 1) were cut from supporting poles of feeding bridges in rearing ponds with shrimp negative for EHP infection (Chantaburi province, Thailand). The mussels were cleaned thoroughly 3 times with clean tap water to remove all organisms attached to the shells. The naïve mussels were acclimatized in clean artificial seawater for 3 days with 100% water exchange daily. Five mussels were dissected out of their shells and samples of the whole body were preserved in lysis buffer for DNA extraction and species identification by PCR using 18S rRNA gene and the PCR products were subjected to sequence analysis (Macrogen, Korea). EHP detection was carried out by EHP-SWP nested PCR analysis. The eukaryotic 18S rRNA gene primers used were UnivF-15 (5’-CTGCCAGTAGTCATATGC-3’) and UnivR-1765 (5’-ACCTTGTTACGACTTTAC-3’) (Frischer et al., 2000). The expected product size was 1,769 bp. The remainder of the cleaned mussels were kept in a stock maintenance tank (100 L-tank, at 28°C) containing 20 ppt artificial sea water with feeding for 7 days before proceeding with EHP cohabitation assays.

**Figure 1.**
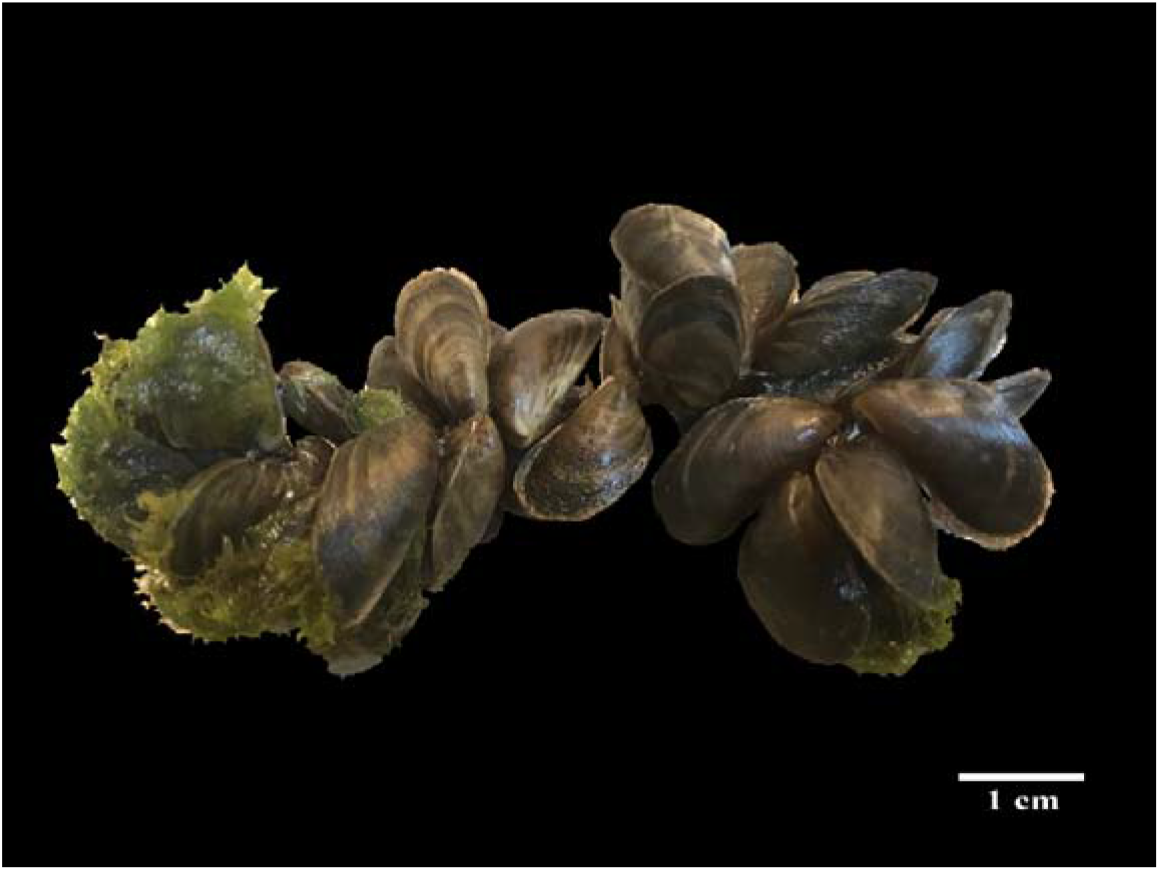
Photograph of the false mussel, *Mytilopsis leucophaeata* collected from the EHP affected ponds.

### Cohabitation bioassay between EHP infected shrimp and naïve mussels

This experiment (first cohabitation test) was designed to determine whether EHP-positive shrimp could transmit EHP to naïve *M. leucophaeata*. Ten EHP-infected shrimp and 150 of EHP-free mussels were co-cultured for 20 days in a 100 liter-plastic tank containing 20 ppt artificial saline water 28°C and sufficient aeration (Dissolved oxygen > 4 ppm). There was neither water recirculation nor water exchange except for refilling with clean saline water to replace any rearing water lost to evaporation. The shrimp were fed 2 times daily with 3% body weight of commercial pelleted feed. On day 7 and day 20, 30 post cohabitation, mussels were removed for testing. Altogether, 10 were tested for EHP load using the nested SWP-PCR method (Jareonlak *et al*., 2016). These were aggressively surface cleaned 3 times using tap-water for external decontamination. Then, the mussel bodies were removed from their shells and washed with sterilized water before PCR testing. In addition to PCR testing, wet mounts (n=5), H&E staining histology, *in situ* hybridization (ISH) (n=10) and transmission electron microscopy (n=5) were carried out to test for EHP infection in the mussels. The remainder of the mussels from this experiment were used in a second cohabitation experiment in which they served as the potential source of EHP.

### Cohabitation assay between EHP-infected mussels and naïve shrimp

This experiment (second cohabitation test) was designed to determine whether EHP-positive *M. leucophaeata* could transmit EHP to naive shrimp. EHP-infected mussels from the previous cohabitation test with EHP-infected shrimp were aggressively cleaned externally before use in the experiment. Sixty EHP-infected mussels remaining from the previous cohabitation test were co-cultured with 10 EHP-free shrimp in a 100 liter-tank for 10 days. This experiment began two days after the end of the previous cohabitation assay (i.e., after the positive PCR test result for EHP was obtained). The shrimp were fed in the same manner as in the first cohabitation bioassay. At day 10 post cohabitation, individual shrimp (n=8) were dissected longitudinally to divide the HP into 2 halves. One half was used for DNA extraction followed by nested SWP-PCR for EHP detection. The other half was investigated by normal histology and ISH.

### Histological examination

Ten mussels and HP from 10 shrimp were individually fixed in Davidson’s AFA fixative (Bell and Lightner, 1988) for 18-24 hours and processed for normal histological analysis as previously described (Bell and Lightner, 1988). Briefly, the fixative solution was replaced with 70% ethanol overnight before processing, embedding, sectioning (4 µm thick) and staining with hematoxylin and eosin (H&E).

For *in situ* hybridization (ISH), a fragment of the EHP small subunit ribosomal RNA (SSU rRNA) gene was used to generate an EHP probe with a PCR-DIG labeling kit (Roche, Germany) ENF779 and ENR779 (Tangprasittipap *et al*., 2013). The specific probe for white spot syndrome virus (WSSV) gene was prepared to serve as a negative control DNA probe using the same DIG-labelling kit with primers 447F and 447R (Srisala et el., 2008). The reason for using a non-specific DNA probe for EHP instead of no probe as a negative control was to eliminate the possibility of obtaining false ISH reactions that might arise from non-specific binding of DNA as sometimes occurs, for example, with chitin in shrimp samples. The sizes of the amplicons for the SSU and WSSV probes were 779 bp and 447 bp, respectively. The PCR products obtained were purified using a Gene pHlow™ Gel/PCR Kit (Geneaid). Adjacent paraffin sections were prepared for H&E staining and for positive and negative control ISH tests as previously described (Tangprasittipap *et al*., 2013).

For TEM analysis (n=5), small pieces (2 mm^3^) of digestive gland tissue from 20 days co-cultured *Mytilopsis* sp. specimens were fixed in 4% glutaraldehyde in 0.1M PB buffer and were proceed following the procedure of Tourtip et al. (2009). The uranyl acetate and lead citrate stained sample sections on copper grid were examined using a Hitachi H-800 transmission electron microscope.

For wet mount analysis, digestive glands of mussels (n=5) were dissected and externally decontaminated using sterilize distilled water before they were sliced into thin pieces. Then, the tissue pieces were placed on glass slide and gently crushed before being staining with 2% phloxine B followed in order to observe viability of EHP spores by polar tube extrusion (Aldama-Cano et al. 2018).

## RESULTS AND DISCUSSION

### Identification of mussels in ponds with EHP-infected shrimp

Of the 21 ponds surveyed, 6 exhibited shrimp with severe EHP infections and also contained what are locally called false mussels. Using the eukaryotic 18S rRNA gene primers with 6 mussel samples arbitrarily collected from such farms yielded amplicons of 1,741 nucleotides and sequencing revealed that all showed 99.94% identity to the GenBank database record KX713323.1 for *M. leucophaeata*. This is the black, false mussel that is a suspension filter feeder in class Bivalvia. Although there are as yet, no confirmed reports of EHP parasitizing this mussel, there have been reports of some samples being PCR positive for EHP. On the other hand, there are many reports of other microsporidians in various tissues of bivalves. Examples include one in the connective tissue of the digestive gland of the scallop *Aequipecten opercularis* (Lohrmann et al. 2000), one in the connective tissue surrounding the gut epithelium of the oyster *Ostrea lutaria* (Jones 1981), and one in the stomach epithelium of clams *Ruditapes decussatus, Venerupis pullastra* and *V. rhomboides* (Villalba et al. 1993a,b).

### Naïve mussels cohabitated with EHP-infected shrimp become PCR positive for EHP

Sampling of 10 arbitrarily selected mussels from the stock batch of 300 mussels collected from a shrimp pond without EHP-infected shrimp revealed that all 10 samples were negative for EHP by the SWP-PCR method. The number of 150 mussels from this batch were arbitrarily selected to cohabitate with 10 shrimp from a pond severely infected with EHP for 7 days. The results showed that 2 out of 10 sampled mussels (20%) gave positive PCR test results for EHP and after 20 days of cultivation, 10/10 sampled mussels (100%) were positive (Fig. 2).

**Figure 2:**
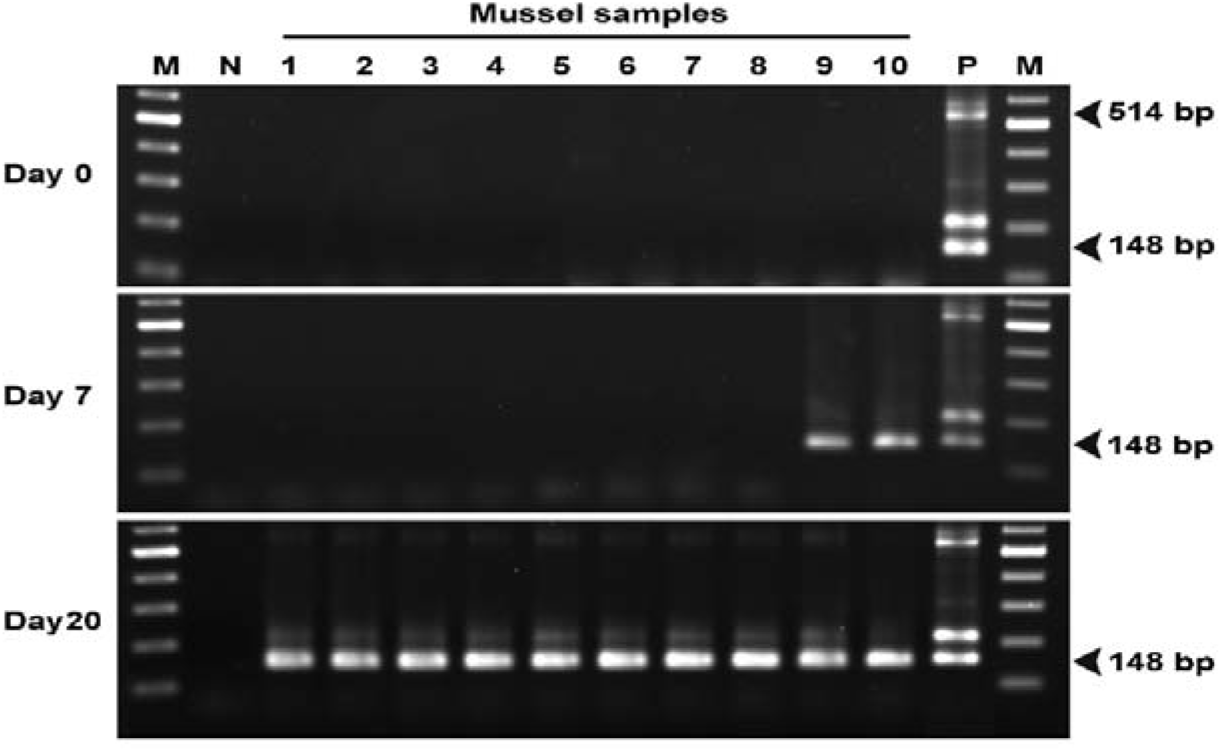
Agarose gel electrophoresis using SWP-PCR detection of the samples collected at day 0, 7 and 20 post cohabitation in the first cohabitation experiment between EHP infected shrimp and EHP-free mussels (*M. leucophaeata*). Lane M: 2 log marker Lane N: negative control using sterile water as template; Lane 1-10: 10 mussels collected each day; Lane P: positive control using DNA extracted from EHP-infected shrimp as template. The expected amplicon sizes for the first step and the 2nd-step SWP-PCR are 514 and 148 bp, respectively.

### EHP transmitted from SWP-PCR positive mussels to cohabitated naïve shrimp

For the second cohabitation test, *M. leucophaeata* from the first cohabitation test were aggressively washed before cohabitation with EHP-negative shrimp (no positives for 5 shrimp pre-sampled by SWP-PCR). At day 10 of cohabitation, 3/8 (37.5%) collected shrimp demonstrated SWP-PCR positive (Fig. 3). This is similar to transmission in shrimp-shrimp cohabitation tests that give relatively low transmission at 7 days but 100% transmission by 14 days (Salachan et al., 2017). Thus, the results were sufficient to reveal that EHP infections could be transmitted from mussels to shrimp.

**Figure 3:**
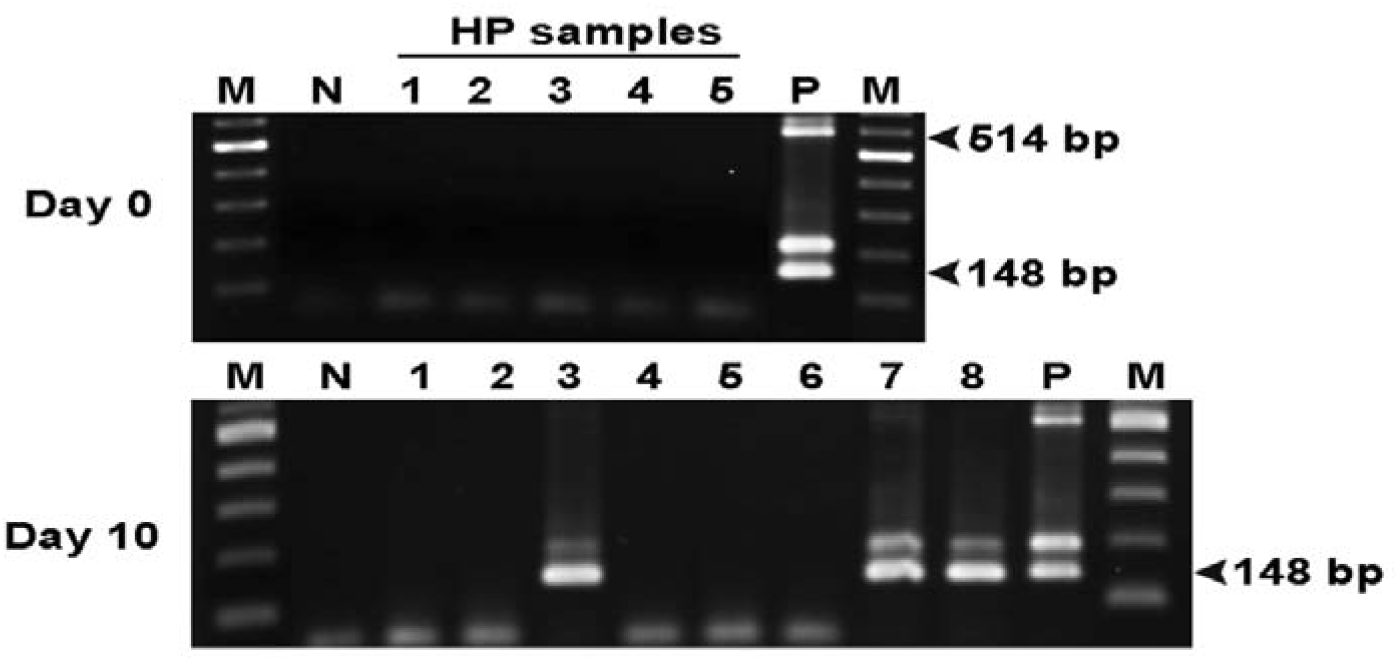
Agarose gel electrophoresis using SWP-PCR detection of the samples collected at day 0 and 10 post cohabitation in the second cohabitation experiment between naïve shrimp and EHP infected *M. leucophaeata*. On day 0 post cohabitation, the number of 5 shrimp were tested (Lane 1-5) and on day 10 post cohabitation, the number of 10 shrimp were tested (Lane 1-10). Lane M: 2 log marker Lane N: negative control using sterile water as template; Lane P: positive control using DNA extracted from EHP-infected shrimp as template. The expected amplicon sizes for the first step and the 2nd-step SWP-PCR are 514 and 148 bp, respectively.

### M. leucophaeata positive for EHP by SWP-PCR shows no signs of EHP infection

At the end of the first cohabitation assay with EHP-infected shrimp, the mussel samples (n=10) examined by normal histological analysis (H&E stained slides) (Fig. 4) showed no signs of EHP infection. However, adjacent tissue sections subjected to ISH using an SSU rRNA probe for EHP gave diffuse positive signals for the presence of EHP-DNA in the cytoplasm of only epithelial cells of the digestive gland (Fig. 5). These ISH signals seen in 4/10 specimens were strongest in the cytoplasm nearest the digestive gland lumen side of the cells. Despite these diffuse ISH reactions, inspection of both the H&E stained slides and slides of adjacent sections stained by ISH showed no indications of microsporidian developmental stages such as plasmodia or spores. Nor were any microsporidian-like structures or positive ISH reactions observed in other tissues (i.e. gills, gonads, intestine and stomach). Since we used a non-specific DNA probe (WSSV) rather than no probe as the negative control, it was very unlikely that the ISH signals arose from non-specific DNA binding as can sometimes be seen with chitin in shrimp.

**Figure 4:**
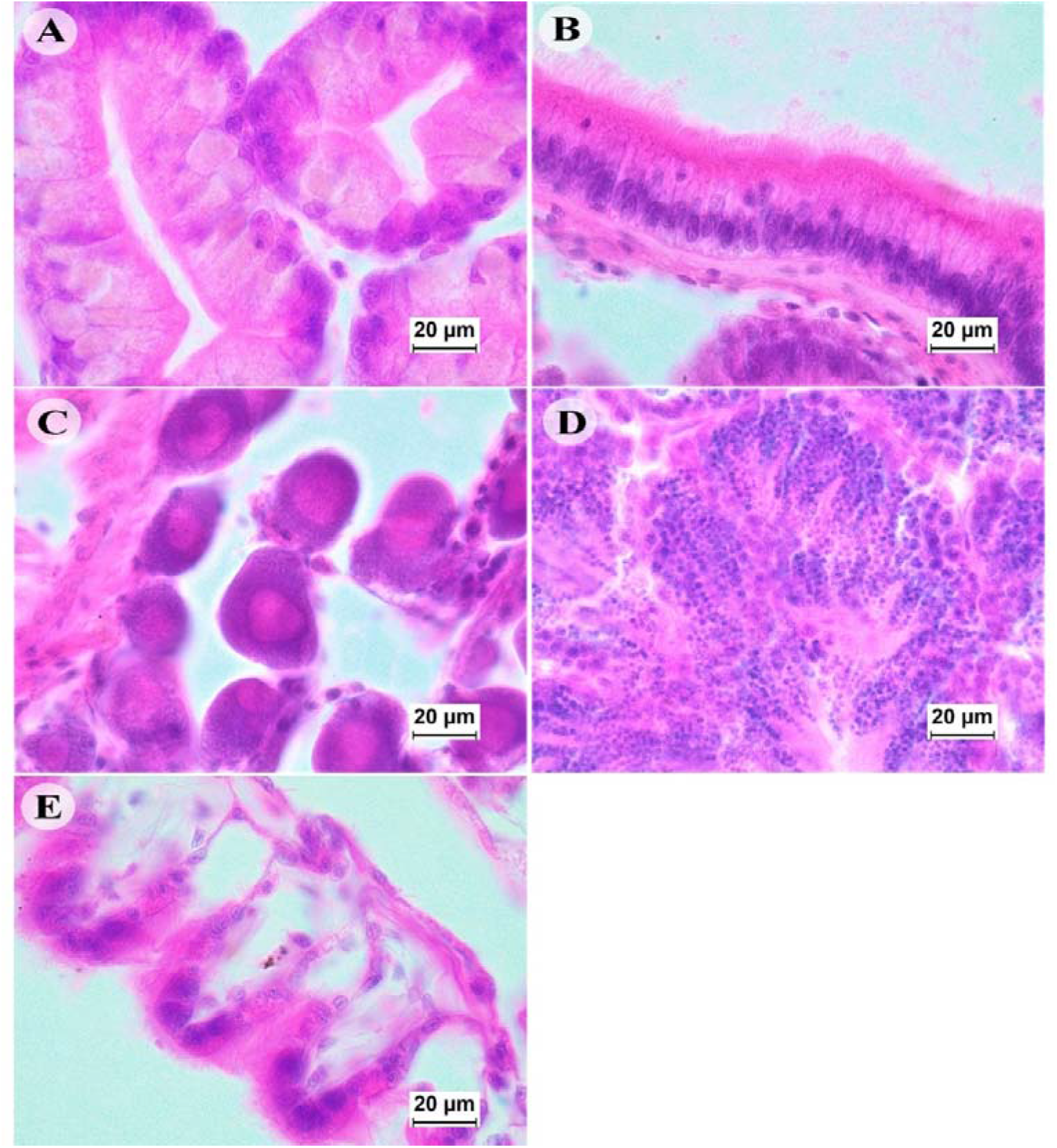
Photomicrograph of histological examination of EHP-PCR positive *M. leucophaeata* revealed the absence of EHP in the mussel tissue including (A): digestive gland, (B): intestine, (C and D): female and male reproductive organ, respectively and (E): gill (H&E staining, 100x magnification).

**Figure 5:**
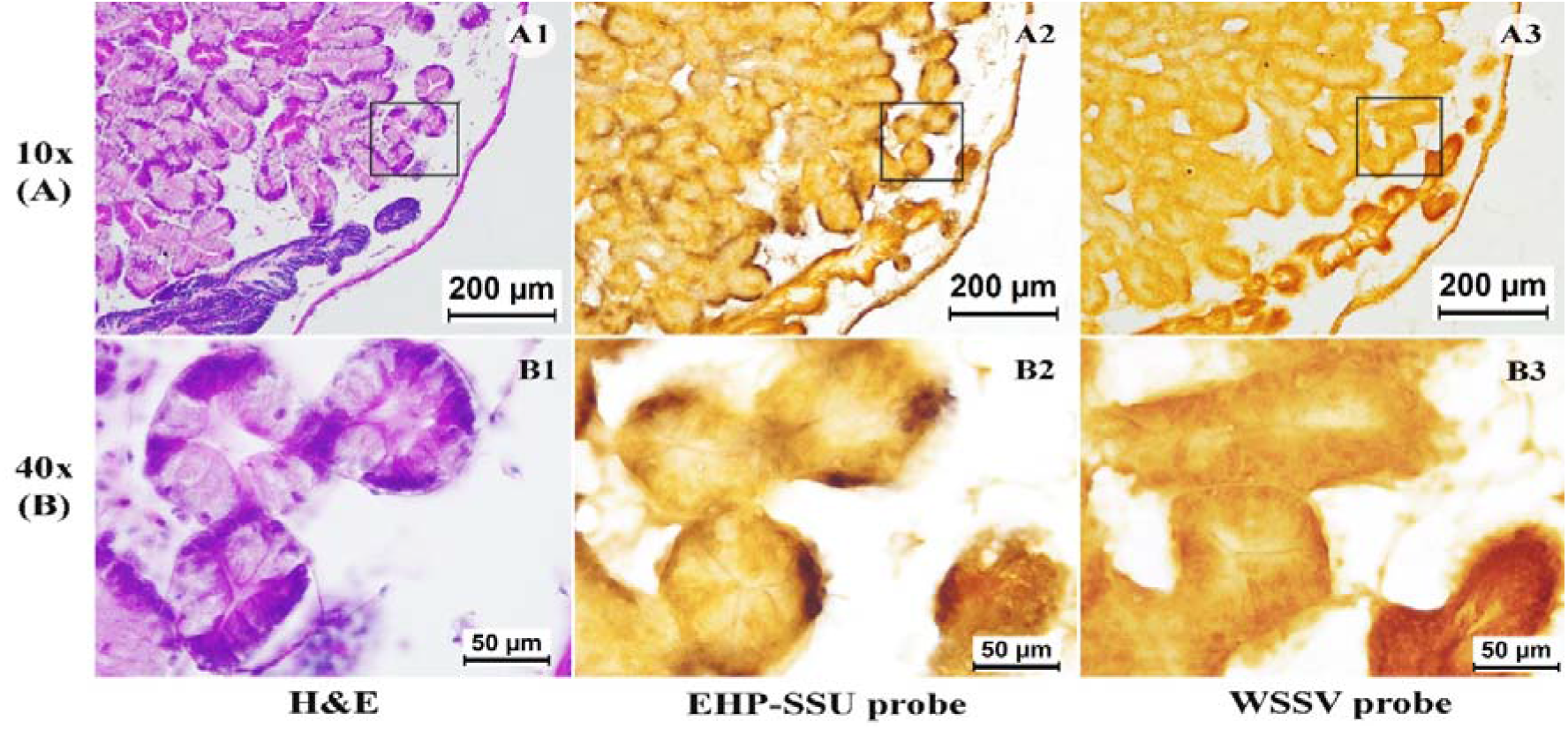
Photomicrographs of H&E staining and ISH of digestive glands of EHP-PCR positive *M. leucophaeata* at 10x (A) and 40x magnification (B). The box area in (A) was magnified in (B). 1-3: the EHP-infected digestive gland stained with H&E, EHP-SSU probe and WSSV probe, respectively. A1 and B1 showed H&E stained digestive gland without any EHP stage inside the cells, A2 and B2 showed the diffuse positive signals of EHP-SSU probe in the cytoplasm of epithelial cells of the digestive gland, and A3 and B3 revealed none of WSSV probe reaction with digestive gland.

For further examination of *M. leucophaeata* from this cohabitation trial, the digestive gland tissues were studied, since some of the histological specimens had given positive ISH results for EHP. As with the histological analysis by light microscopy, none of the epithelial cells of the digestive gland revealed the presence of microspridian-like life stages (e.g., sporoplasts, plasmodia or spores) by TEM (Fig. 6). On the other hand, wet mount analysis of the digestive gland with phloxine-B staining revealed the presence of EHP-like spores, some of which had extruded polar tubes (Fig. 7). Polar tube extrusion upon phloxine-B staining is used as an indication of viable spores (Aldama-Cano et al. 2018).

**Figure 6:**
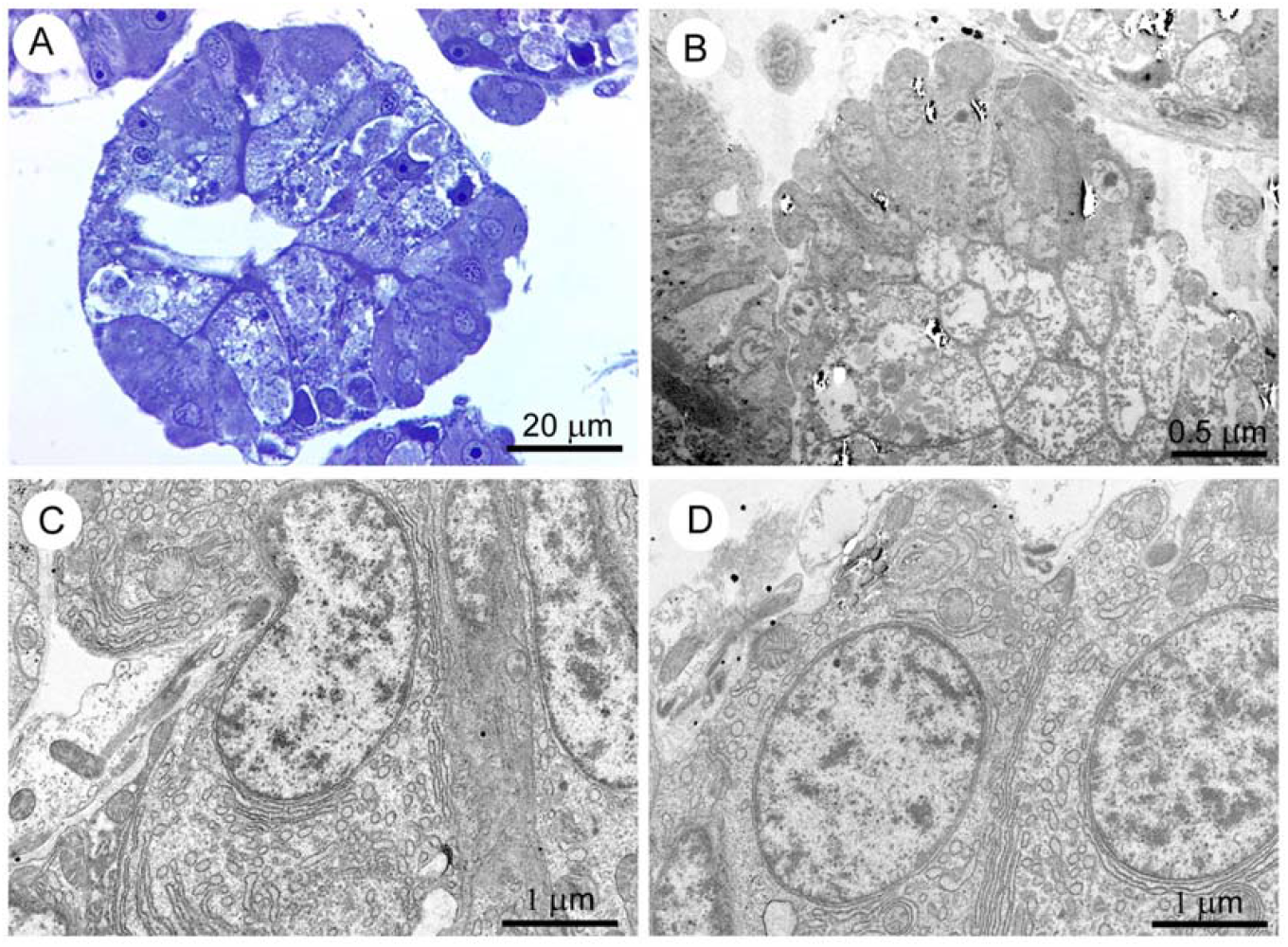
Photomicrographs of semithin (1 µm-thick; A) for light microscopic examination and thin (∼70 nm-thick; B-D) sections for transmission electron microscopic examination of mussel digestive glands. A and B: The epithelial cells of digestive glands tested positive for EHP infection by ISH using EHP-SSU probe showed the absence of intracellular life stage of EHP. C and D: Transmission electron micrographs of the epithelial cells showed euchromatic nucleus and cytoplasm containing numerous rough endoplasmic reticulum and a few mitochondria with no EHP.

**Figure 7:**
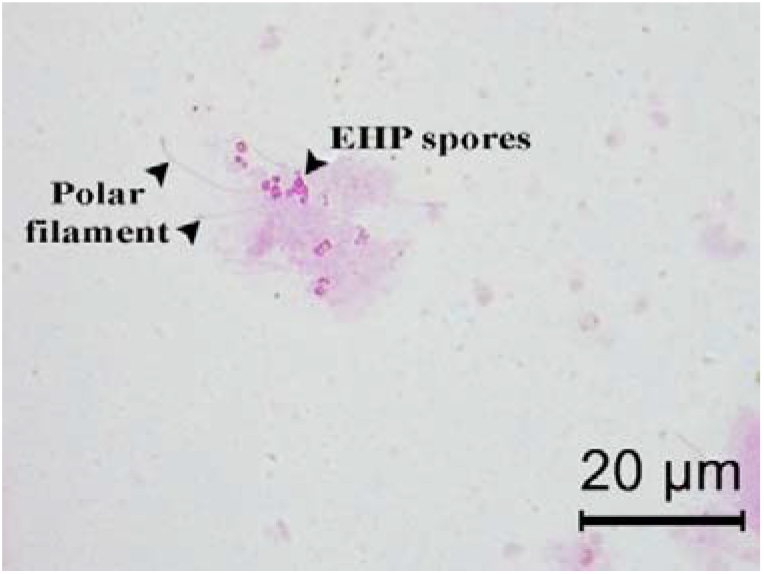
Photomicrograph of wet mount analysis of digestive gland demonstrated the presence of the oval shape-EHP spores and their polar tube extrusion by phloxine B staining (100x magnification).

For the second cohabitation assay employing EHP-positive mussels with naïve shrimp, examination of the shrimp at Day 10 of cohabitation by light microscopy with H&E stained slides clearly revealed EHP spores and plasmodia in the hepatopancreatic tubule epithelium by (Fig. 8, D and E). Adjacent sections stained for ISH gave positive signals for EHP in the cytoplasm of HP epithelium containing microsporidian developmental stages (Fig. 8, B). These results not only confirmed the presence of EHP but also served as a positive control for the EHP ISH-probe, the specificity of which was confirmed by the absence of any ISH signal in the adjacent, negative control tissue section probed with the WSSV-DNA probe (i.e.. the positive ISH signals in the shrimp and mussel tissues could not have arisen from non-specific binding of DNA).

**Figure 8:**
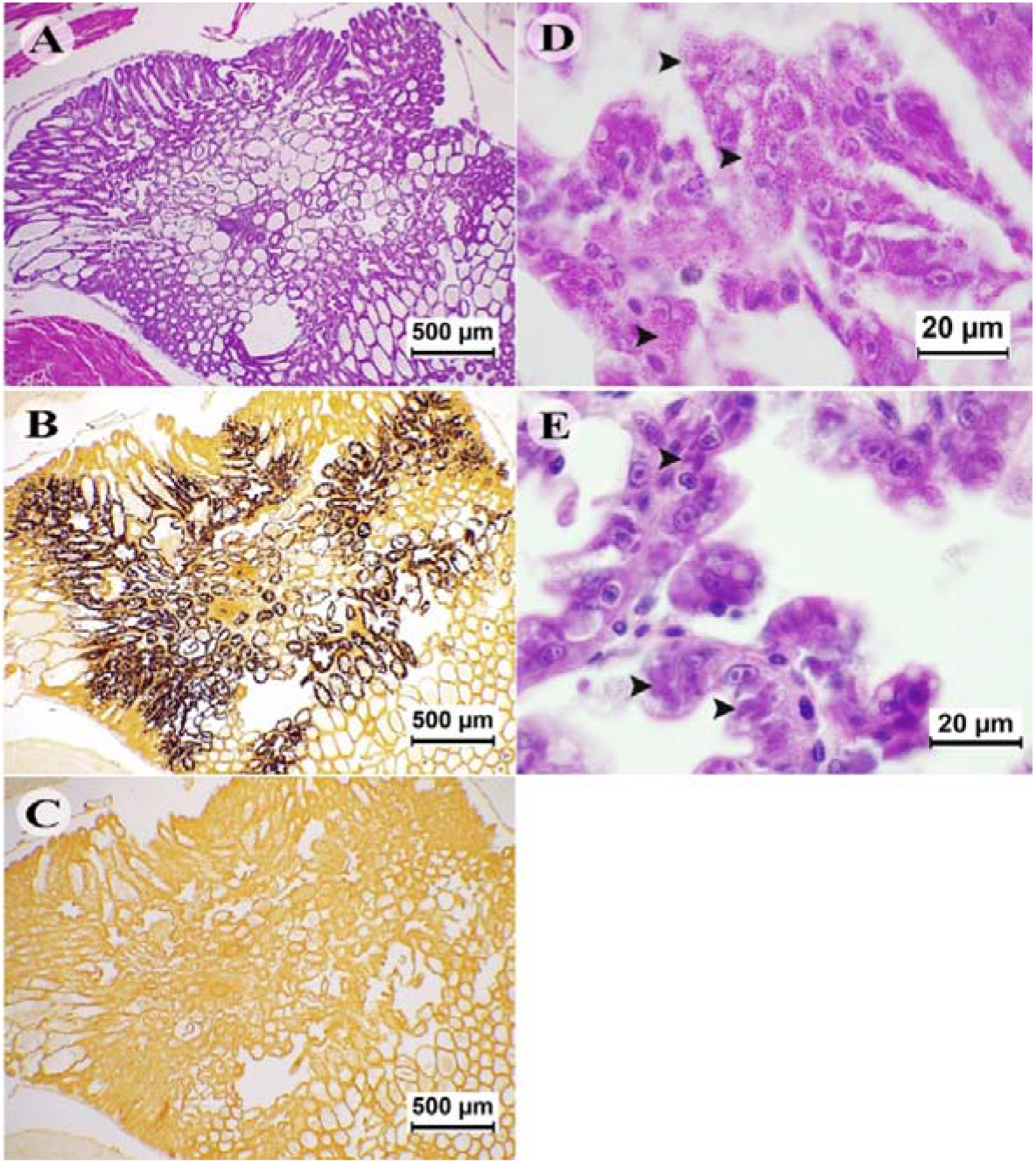
Photomicrograph of EHP infected hepatopancreas of *P. vannamei* stained with H&E and ISH. The adjacent sections of shrimp tissue stained with H&E (A), with the EHP-SSU probe and WSSV probe (B and C). The extensive infection is revealed by *in situ* hybridization (dark brown to black staining, B). Higher magnification (100X) of the hepatopancreatic cells of the EHP-infected shrimp obviously showed structure of EHP spore (arrow, D) and plasmodium (arrow, E).

The two cohabitation assays indicated that *M. leucophaeata* can carry infectious spores of EHP after being co-cultured with EHP-infected shrimp for 7 days and that the spores they carry can be infectious for shrimp. This was confirmed by the observation that active EHP spores could be found in the mussel digestive gland from which it they had not been precluded from it in pseudo-feces. Thus, they could possibly serve as a portion of the mussel diet and could potentially expose the mussels to EHP infection if they were susceptible. Indeed, microsporidian infections have been described in digestive organs of other bivalves such as scallops (i.e. *Aequipecten opercularis*) (Lohrmann et al. 2000). However, we could not find any signs of microsporidian stages in the digestive glands of our EHP-positive mussels by light or electron microscopy. The presence of these spores could have given the SWP-PCR positive results we obtained from the mussel tissues. At the same time, assuming that the EHP spores were selected as a part of their diet, it is possible that the diffuse positive ISH reactions we saw in the digestive gland epithelial cells arose from incompletely degraded DNA of ingested EHP spores.

This proposal is consistent with the knowledge that digestive glands of bivalves contain 2 types of cells, basophilic and digestive cells. The basophilic cells mainly function to produce and secrete enzymes (Dimitriadis et al., 2004) that might include chitinases since they have been reported from *Dreissena*, a sister genus of *Mytilopsis* in the family *Dreissenidae* (McCartney et al., 2019). The digestive cells are known to play an important role in absorption of digested or partially digested nutrients, in intracellular digestion and in antioxidant defense (Faggio et al., 2016). Thus, it is possible that the diffuse ISH signals we saw in the digestive gland arose from partially digested EHP-DNA that had been absorbed by the cells of the digestive gland but still remained sufficiently intact to hybridize with the SSU rRNA gene probe.

## CONCLUSIONS

Our results revealed that *M. leucophaeata* can accumulate EHP spores that are released from EHP-infected shrimp but that they do not become infected and so cannot amplify EHP. However, if removed from a pond containing EHP-infected shrimp and moved directly to an uninfected shrimp pond, they can serve as passive carriers to transmit infective spores to naïve shrimp. Similarly, mussels would constitute a risky live feed for EHP-free broodstock in a shrimp hatchery if not depurated, frozen or pasteurized as previously recommended. On the other hand, since the mussels take up EHP spores (presumedly as food), it may be worthwhile to test banks of them for efficacy in EHP spore removal from contaminated water. If efficacious, they might serve as active biological filters for EHP-spore removal.

## ACKNOWLEDGEMENT

This work was supported by grant from Newton fund under the project “Newton Institutional Links 2017 and Newton Prize (Chairman awards 2019)” to Prof. Grant D. Stentiford (Cefas/UK) and Dr. Kallaya Sritunyalucksana (BIOTEC, NSTDA/Thailand). A Post-doctoral fellowship granted to Dr. Natthinee Munkongwongsiri was from the National Center for Genetic Engineering and Biotechnology (BIOTEC), a member of National Science and Technology Development Agency (NSTDA), Thailand. We are very thankful to Juree farm for all facilities supporting during shrimp and *M. Leucophaeata* collection in this study. We would also like to thank Prof. T.W. Flegel for assistance in reviewing and editing the drafts of the manuscript.

